# Inhibition of pyruvate dehydrogenase affects the brain protein acylation stronger than PDHA phosphorylation at Ser293

**DOI:** 10.1101/2022.11.10.515938

**Authors:** Vasily A. Aleshin, Daria A. Sibiryakina, Alexey V. Kazantsev, Anastasia V. Graf, Victoria I. Bunik

## Abstract

Adaptation of an organism to metabolic challenges requires mechanisms coupling metabolism to gene expression. Acylations of metabolic and histone proteins acquire significant attention in this regard. We hypothesize that adaptive response to inhibition of a key metabolic process, catalyzed by the acetyl-CoA-generating pyruvate dehydrogenase (PDH) complex, may be mediated by changed protein acylations. The hypothesis is tested by intranasal administration to animals of PDH-specific inhibitors acetylmethylphosphinate (AcMeP) or methyl ester of acetylphosphonate (AcPMe), followed by assessment of physiological parameters, brain protein acylation system and expression/phosphorylation of PDHA subunit. At a fixed dose, AcMeP, but not AcPMe, decreases acetylation and increases succinylation of the brain proteins of apparent molecular mass of 15-20 kDa. Regarding the 30-50 kDa proteins, a strong inhibitor AcMeP affects acetylation only, while a less efficient AcPMe mostly increases succinylation. No increase in the succinylation of the 30-50 kDa proteins by AcMeP coincides with its induction of desuccinylase SIRT5, not observed in the AcPMe-treated animals. The brain PDHA expression or phosphorylation, the animal behavior or ECG do not significantly differ between the studied animal groups. The data indicate that a short-term inhibition of the brain PDH affects acetylation and/or succinylation of the brain proteins, dependent on the inhibitor potency, protein molecular mass and acylation type. Homeostatic nature of these changes is implied by stability of physiological parameters after the PDH inhibition.

## INTRODUCTION

Protein acylation has long been known to regulate the chromatin organization by histones [1–3] and the function of 2-oxo acid dehydrogenases, whose multienzyme complexes produce activated acyl residues in the form of the enzyme intermediates and acyl-CoA products [4,5]. Recently, post-translational modification of the protein lysine residues by acylation has acquired increased attention regarding non-nuclear proteins beyond 2-oxo acid dehydrogenases, whose deacylation is catalyzed by “rejuvenating” factors, NAD+-dependent sirtuins [6–8]. At the same time, nuclear localization of 2-oxo acid dehydrogenase complexes and their participation in histone acylation have been revealed [9–12], as well as the role of mitochondrially localized 2-oxo acid dehydrogenase complexes in acylation of variety of metabolic proteins [13,14]. Thus, production of the acylating residues by 2-oxo acid dehydrogenase complexes may provide acyl-CoAs for acylation of histone and non-histone proteins. As the process is tightly coupled to the brain glucose metabolism where these complexes occupy a key position, the regulatory role of the protein acylation may be involved into homeostatic responses to metabolic perturbations. Indeed, this is revealed in a recent study of the seizures-induced disbalance of the brain metabolism in a rat model of epilepsy [15].

On the other hand, acyl-CoAs may also arise from other metabolic pathways, such as fatty acid oxidation, or metabolism of ketone bodies. In particular, in specific experimental systems, acetate-dependent acetyl-CoA synthetase 2 (ACSS2) or citrate-dependent ATP-citrate lyase may be involved in histone acetylation [16,17]. Worth noting, the relative substrate fluxes through the acyl-CoA-generating pathways depend on the tissue-specific metabolism and protein expression. Besides, different physiological or pathological conditions may change contributions of specific metabolic fluxes to protein acylation. For instance, acetyl-CoA generated by fatty acid oxidation drives excessive hepatic acetylation in mice [18]. When cultured human fibroblasts are stimulated to oxidize fatty acids, acetylation of proteins also increases [18]. However, such stimulation or protein hyperacetylation correspond to pathological conditions. Physiological level of the protein acetylation may well be contributed by other acetyl-CoA sources, such as PDH.

Acylation of brain proteins is of special interest in view of molecular mechanisms of memory and cognition. Histone acetylation is essential for regulation of memory storage, with learning and memory requiring the chromatin remodelling, dependent on histone acetylation [19]. A long-known decrease in the activity of 2-oxoglutarate dehydrogenase (OGDH) complex in the Alzheimer’s disease (AD) brains [20] may underlie a recently revealed decrease in succinylation of mitochondrial proteins in these brains [21].

The goal of this work is to characterize an impact of PDH-dependent acetyl-CoA production on acylation of the brain cortex proteins in the rat model where the brain responds to a short-term metabolic challenge. The challenge may arise upon environmental or similar stress to be addressed by physiological mechanisms of homeostasis. A short-term reversible PDH inhibition by pharmacological means is a better mimic of such stresses than manipulation of gene expression or protein translation, both inducing secondary effects associated with impaired protein-protein interactions network [22–25]. Based on previous work on catalysis-dependent selectivity of the inhibitory action of the phosphonate and phosphinate analogs of pyruvate in vivo [22,23], and up to a 100-fold higher efficacy of the latter vs former [23–25], we employ the two synthetic analogs of pyruvate, shown in Figure 1. Having similar electrostatic and steric properties, AcMeP and AcPMe should not significantly differ in their bioavailability. Hence, administration of the same dose of these inhibitors that strongly differ in their inhibitory power [23–25] models increasing levels of PDH inhibition in the brain. In view of the recently characterized interdependency of the different types of the protein acylations due to the overlapping sites and competition between the PDH and OGDH complexes for their common substrate CoA [15], we assume that PDH inhibition may not only have a direct effect on the protein acetylation, but also indirectly affect other types of acylations. From those, succinylation may be the most affected one, as there is the tight metabolic connection between PDH and OGDH through the tricarboxylic acid (TCA) cycle, where PDH complex feeds acetyl-CoA, while the succinyl-CoA producer, i.e. OGDH complex, limits the substrate flux [25,26]. Thus, the brain response to PDH inhibition is characterized in this work by study of both the acetylation and succinylation of the brain proteins.

**Figure 1.**
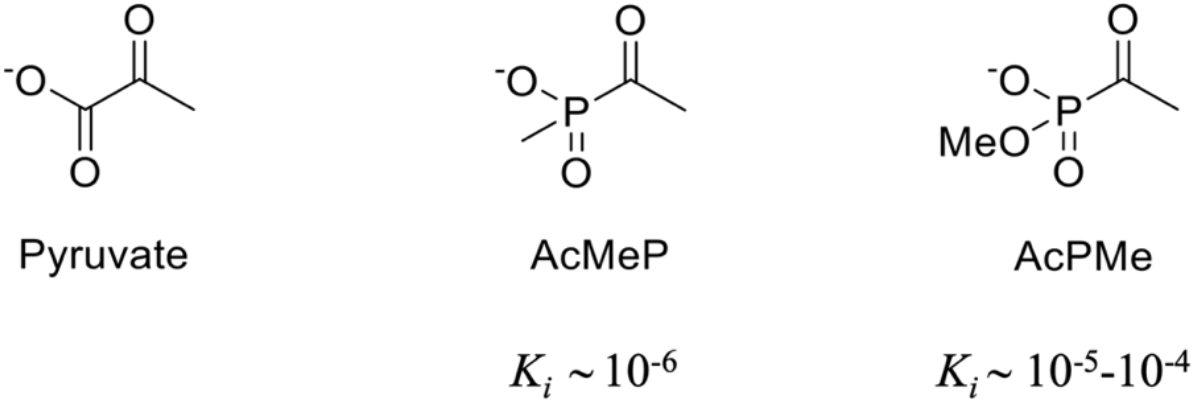
Structures of the pyruvate dehydrogenase 2-oxo substrate (pyruvate) and its phosph(i/o)nate analogues acetylmethylphosphinate (AcMeP) and methylated ester of acetylphosphonate (AcPMe). Inhibition constants (Ki) of the phosphinate and phosphonate analogs are from the published results [23–25].

## MATERIALS AND METHODS

### Reagents

Sodium salts of AcPMe and AcMeP were synthesized as published before [24] and their purity was checked by NMR. The buffers, salts and other reagents were of the highest purity available and obtained from Merck (Helicon, Moscow, Russia). Deionized MQ-grade water was used for solution preparations. The used antibodies are indicated in section 2.7.

### Animal Husbandry

Animal experiments were performed according to the Guide for the Care and Use of Laboratory Animals published by the European Union Directives 86/609/EEC and 2010/63/EU, and were approved by the by Bioethics Committee of Lomonosov Moscow State University (protocol number 139-a from 11 November 2021).

Wistar male rats were purchased from the Russian Federation State Research Center Institute of Biomedical Problems RAS (IBMP). Rats of 11–12 weeks old (weighting 320 ± 30 g) were used in the experiment. Animals were kept at 21 ± 2°C and relative humidity 53 ± 5% with the 12/12 h light/dark cycle (lights on 9:00 = ZT 0, lights off 21:00 = ZT 12). They had free access to standard rodent pellet food and tap water. Each cage contained 4-5 animals, standard T/4K cages (555/4K, 580×375×200 mm) were used.

### Administration of the Pyruvate Dehydrogenase Inhibitors

Intranasal administration of a water solution of AcPMe or AcMeP (at a dose of 0.1 mmol/kg) was performed once in the morning at 10:00 ± 1 h. The animals were randomly divided into groups, one receiving the water solution of AcPMe (n=8), the second - the water solution of AcMeP (n=8) and the third – the physiological solution, i.e. 0.9% NaCl (n=6). A total of 22 rats were used in the work. Intranasal application was used as a non-invasive method providing an access to the central nervous system for different molecules, including those which do not cross the blood-brain barrier.

A day (24 h) after the administration of the inhibitors, the physiological tests were made (described in chapter 2.4), followed by decapitation using a guillotine (OpenScience, Russia). Euthanasia by decapitation is considered one of the least stressful methods to kill young rats [27–29]. The guillotine was washed thoroughly with water and ethanol after each use, so that no blood smell could stress another rat. The procedure was done according to recommended protocols [27–29]. The extracted brains were immediately transferred to ice. The cerebral cortex (cerebral hemispheres) was separated and frozen in liquid nitrogen 60–90 s after decapitation. The cortices were stored at −70◦C before biochemical analyses.

### Physiological Tests

Estimation of behavioral activity was performed in the “Open Field” test as described before [30]. A circular polypropylene arena («OpenScience», Moscow, Russia) was used for testing. The estimated parameters (duration and number of grooming acts, duration of freezing, number of rearing acts and number of crossings for any type of lines) served as indicators of anxiety, exploratory and locomotor activity according to established views [31,32].

ECG was recorded on awake animals within the registration time of 3 min using non-invasive electrode placement as previously described [30]. Briefly, the following parameters were assessed through ECG: the average interval between the heartbeats (R-R interval, ms), the standard deviation of the average R-R interval values (SD, ms), the range of R-R interval values, i.e., the difference between the maximal and minimal values (dX, ms), the root mean square of successive differences in R-R intervals (RMSSD, ms), and stress index (SI, arbitrary units).

### Preparation of Homogenates of the Rat Cerebral Cortex

Homogenization of the tissue and sonication of homogenates was carried out according to the previously published protocol [33].

### Enzyme Activity Measurements

The activity of malate dehydrogenase (MDH) was measured as previously described [34].

### Western-Blotting Quantification of the Protein Levels and Their Post-translational Modifications in Rat Cerebral Cortex

The protein levels of SIRT3, SIRT5, PDHA (alpha subunit of PDH), phosphorylated at Ser293 PDHA and acetylated proteins were estimated by western-blotting using primary antibodies from Cell Signaling Technology (Danvers, MA, United States) with the product numbers #5490, #8782, #3205, #37115 and #9814, respectively. The succinylated proteins were assessed using antibodies from PTM Biolabs (Chicago, IL, United States) #PTM-401. All the primary antibodies were used in 1:2,000 dilutions, with the secondary anti-rabbit HRP-conjugated antibodies from Imtek (Moscow, Russia), #’P-GAR Iss’. The relative quantification of chemiluminescence was performed in ChemiDoc Imager (Bio-Rad, Hercules, CA, United States) and Image Lab software version 6.0.1 (Bio-Rad, Hercules, CA, United States). Normalization of the immunostaining levels to the total protein in the corresponding gel lanes was performed using the protein fluorescent quantification with 2,2,2-trichloroethanol, similarly to the published procedure [35]. Total intensity was determined as integral intensity of the whole track up to 100 kDa.

### Statistics and Data Analysis

Data were analyzed using the GraphPad Prism 7.0 software (GraphPad Software Inc., San Diego, CA, USA). All the data on the graphs were shown as columns with the means and standard errors of mean (SEM), added by the individual values of the parameters for each animal. Associations between the parameters were analyzed using Spearman’s correlations. Statistical significance of differences upon comparison of the three experimental groups was assessed using one-way analysis of variance (ANOVA) with Tukey’s post hoc test. The statistically significant (p ≤ 0.05) differences are shown on the graphs. Significant p-values of the ANOVA factor and the F-values with the degrees of freedom, i.e. F (DFn, DFd), are shown below the graphs. In all these cases, the F-values were larger than the corresponding critical values, supporting the significance of the reported results.

## RESULTS

### AcMeP decreases acetylation of the major protein band of 15 kDa, inducing diverse effects on acetylation of the brain proteins of 30-50 kDa

Western blot with antibodies on acetylated proteins reveals a major band of acetylated brain proteins in the region of 15 kDa (Figure 2A). Significant relative intensity of this band and similarity of its apparent molecular mass to that of core histones [3] suggests that the band comprises the brain histone proteins. The stronger the PDH inhibitor, the more pronounced the effects on the brain proteins acetylation. Indeed, the AcMeP-treated animals show changed acetylation levels in all the four quantified proteins bands, while the treatment with a less efficient inhibitor AcPMe changes acetylation only in the band of 31 kDa (Figure 2, the upper part). A remarkable feature of the AcMeP-induced changes is the diversity of its effects on the acetylation of the brain proteins of different molecular mass. That is, the acetylation of the proteins of 15, 29, 31 kDa is decreased after AcMeP treatment, while acetylation of the proteins of 50 kDa is increased (Figure 2, the upper part). These opposite effects of AcMeP, as well as a low contribution of the band of 31 kDa to the total acetylation, result in no significant change in the total acetylation of the brain proteins after the treatments with either AcMeP or AcPMe.

**Figure 2.**
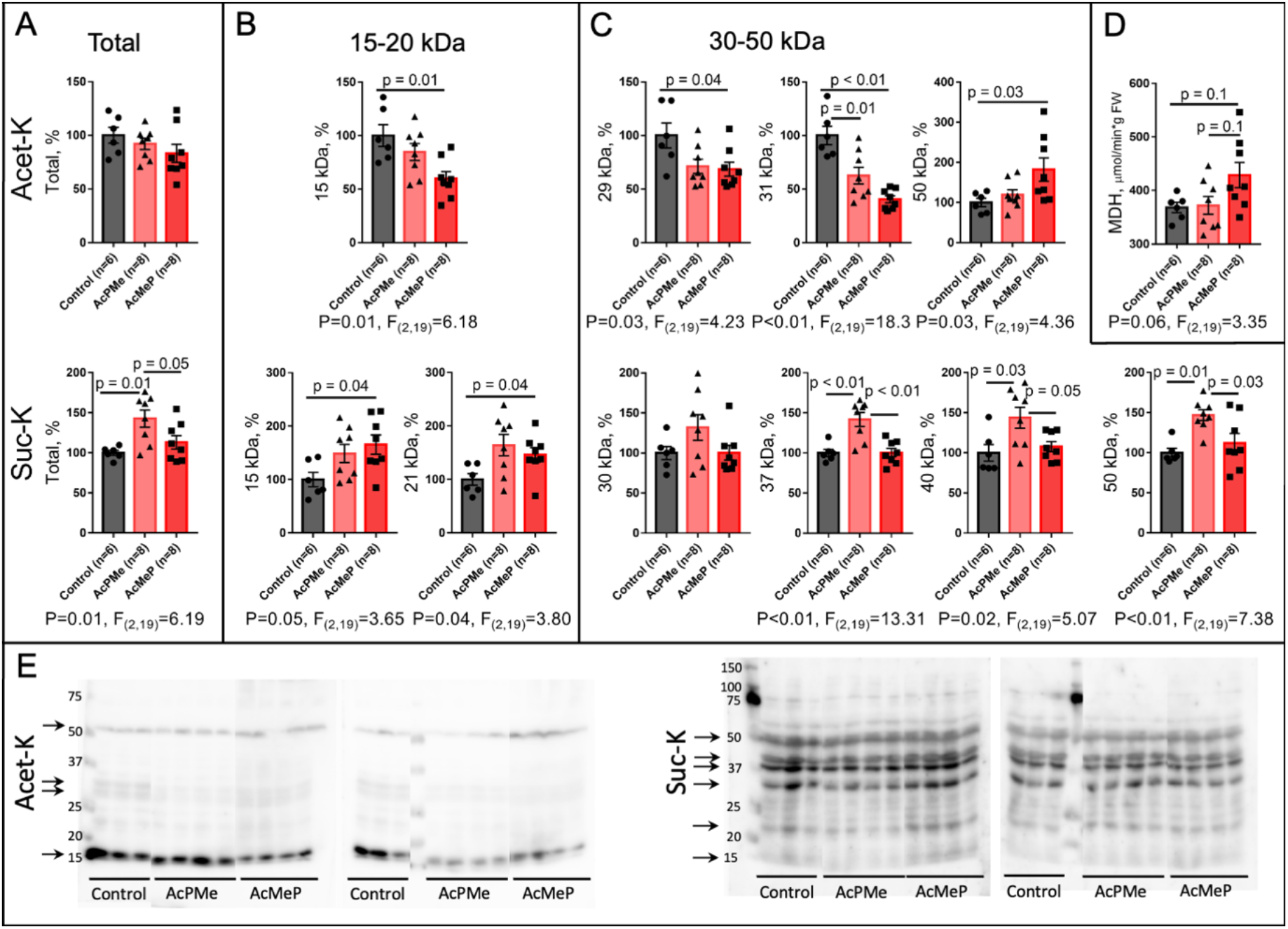
The levels of acylations of diverse proteins in the cerebral cortex homogenates 24 h after administration of PDH inhibitors. The acetylation and succinylation are shown in the upper and lower parts of the figures A-C. Each band intensity is normalized to the total protein in the lane and related to an average value of the normalized band intensity in the control group of rats as described in “Materials and methods”. Significance of the differences between the groups in A-D are indicated by the lines with their p-values above the lines. When the treatment factor is significant according to ANOVA analysis, the characteristic p and F values are given under the graphs. A – Total protein acetylation and succinylation. B – Changes in acetylation of low molecular mass proteins (15-20 kDa). C – Changes in acetylation of the proteins 30-50 kDa. D – Effect of the PDH inhibitors on total activity of malate dehydrogenase (MDH) in the cortex homogenates. E – Western blots of the brain acetylated (left) and succinylated (right) proteins, used for the quantifications presented in A-C. The molecular mass markers are given at the left. The quantified bands are indicated by arrows. Here and further, one panel of the samples combines the relevant parts of one blot.

The AcMeP-induced increase in acetylation of metabolic proteins, determined by Western blot, is independently supported by functional assays. The activity of MDH (36 kDa), known to be elevated upon the protein acetylation [36], shows a trend (p=0.1) to increase along with the AcMeP-induced increase in acetylation of proteins of 50 kDa (Figure 2D). The images of Western blots supporting these graphs are shown in Figure 2E.

### Response of the brain protein succinylation to PDH inhibition is different from that of acetylation3

In contrast to a decrease in acetylation of 15 kDa proteins by AcMeP, the inhibitor increases succinylation of the brain proteins of 15-20 kDa. Once again, the less efficient inhibitor AcPMe does not affect the succinylation of these brain proteins of low molecular mass (Figure 2A-C, the lower part).

However, an opposite situation is observed for succinylation of the brain proteins of 30-50 kDa, that is significantly increased across all the protein bands by the less efficient inhibitor of PDH AcPMe, but is not changed by the strong inhibitor AcMeP (Figure 2A-C, the lower part). As the proteins of 30-50 kDa represent the major part of the succinylated proteins, the total succinylation is also increased by AcPMe, in contrast to AcMeP.

Thus, opposite effects of a strong PDH inhibitor AcMeP are revealed not only regarding the proteins of different molecular masses, but also regarding the different types of the acylations. The AcMeP-induced decrease in the acetylation of proteins with apparent molecular mass close to that of core histones (15-20 kDa) is accompanied by increased succinylation of these proteins, while the succinylation of metabolic proteins is not affected by AcMeP.

### Responses of the mitochondrial deacetylase SIRT3 and desuccinylase SIRT5 to PDH inhibition

In view of the changes in the brain protein acetylation and succinylation by the PDH inhibitors (Figure 2), we assessed the brain levels of the main corresponding deacylases. The changes in the protein expression of the mitochondrial deacetylase sirtuin 3 (SIRT3) and desuccinylase sirtuin 5 (SIRT5) are shown in Figure 3. While no significant changes in the levels of either SIRT3 or SIRT5 are induced by PDH inhibitors, compared to the non-treated animals, the level of SIRT5 exhibits a significant increase after the treatment with AcMeP, compared to that with AcPMe. The AcMeP- vs AcPMe-induced increase in SIRT5 (Figure 3) is in good accordance with the decreased succinylation of the proteins of 30-50 kDa in the AcMeP vs AcPMe animals (Figure 2C, the lower part). The finding is additionally supported by existence of the negative correlations between succinylation of the brain proteins and levels of the desuccinylase SIRT5 in the brain (Table 1). In contrast, protein acetylation levels do not negatively correlate with the levels of mitochondrial deacetylase SIRT3. Instead, total acetylation and acetylation of 15 kDa proteins are positively correlated with SIRT3 levels (Table 1). On the other hand, succinylation of 30 kDa proteins positively correlates with the brain levels of deacetylase SIRT3 (Table 1).

**Table 1.** Spearman correlations of the levels of acylations of diverse proteins (acetylation and succinylation) with those of SIRT3 (deacetylase) and SIRT5 (desuccinylase), determined across the studied animal groups (n=22). Each cell presents the corresponding correlation coefficient (Rs) and significance of the correlation (p). Positive or negative correlations with p < 0.1 are colored in the shades of red or blue, respectively, with statistically significant (p < 0.05) correlations brighter than the trends (0.05 < p < 0.1).

|  | Sirt3 |  | Sirt5 |  |
| --- | --- | --- | --- | --- |
|  | Rs | p | Rs | p |
| Ac-K Total | 0.45 | 0.04 | 0.15 | 0.51 |
| Ac-K 15 kDa | 0.55 | 0.01 | 0.03 | 0.90 |
| Ac-K 29 kDa | -0.02 | 0.92 | -0.26 | 0.24 |
| Ac-K 31 kDa | -0.11 | 0.63 | -0.35 | 0.11 |
| Ac-K 50 kDa | -0.02 | 0.92 | 0.21 | 0.34 |
| Suc-K Total | 0.12 | 0.6 | -0.33 | 0.13 |
| Suc-K 50 kDa | -0.03 | 0.89 | -0.5 | 0.02 |
| Suc-K 40 kDa | 0.30 | 0.18 | -0.37 | 0.09 |
| Suc-K 37 kDa | 0.20 | 0.36 | -0.67 | 0.00 |
| Suc-K 30 kDa | 0.39 | 0.07 | -0.21 | 0.34 |
| Suc-K 21 kDa | 0.11 | 0.62 | 0.05 | 0.84 |
| Suc-K 15 kDa | -0.18 | 0.43 | 0.17 | 0.44 |

**Figure 3.**
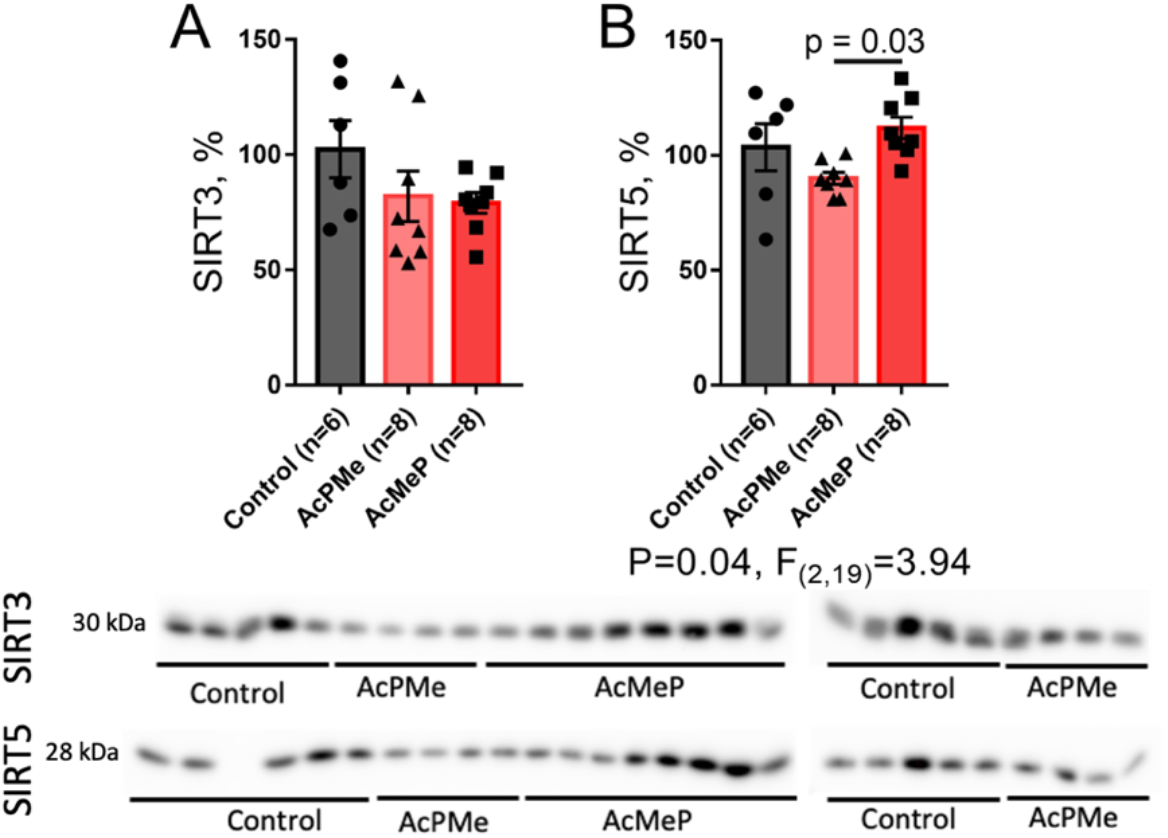
The protein levels of deacylases SIRT3 and SIRT5 upon administration of PDH inhibitors in the cerebral cortex homogenates. A – mitochondrial deacetylase SIRT3; B – deacylase of negatively charged acyl groups SIRT5. Each band intensity is normalized to total protein in the lane and related to an average value of the normalized band intensity in the control group of rats as described in “Materials and methods”. Western blots used for the quantifications are shown below the graphs. Significance of the group differences in the graphs is indicated by lines with their p-values above each line. When the treatment factor is significant according to ANOVA analysis, the characteristic p and F values are given under the graphs.

Thus, the relations between the protein acylations and expression of deacylases may be complex due to potential contribution of multiple factors.

### PDH regulation by phosphorylation of Ser293 of PDHA subunit does not strongly respond to PDH inhibition

Physiological levels of the PDH activity and the enzyme response to multiple factors are controlled by the protein phosphorylation, with the PDH kinase inactivating the enzyme, while PDH phosphatase increasing the activity. Hence, we assessed if the PDH regulation by phosphorylation, as well as the enzyme expression, are involved in the brain response to PDH inhibitors. Figure 4 shows that the brain expression and phosphorylation of Ser293 of PDHA subunit do not strongly change after the administration of the PDH inhibitors. ANOVA analysis reveals a slight increase in the phosphorylated Ser293 after the action of the less effective inhibitor AcPMe (Figure 4B), that may correspond to a more consistent increase in the PDHA expression in this group vs the AcMeP-treated one (Figure 4A). Indeed, the level of the PDHA phosphorylation remains constant, as seen from the ratio of the phosphorylated PDHA to its expression (Figure 4C). Thus, the brain protein acylation responds to PDH inhibition (Figures 2, 3) stronger than the PDH regulation by phosphorylation (Figure 4).

Correlation analysis of the brain protein acetylation levels indicates that expression of the rate-limiting component of the PDH complex, PDHA, is positively correlated with acetylation of 50 kDa proteins, while the acetylation of proteins about 30 kDa is negatively correlated with PDHA expression (Table 2). This finding points to the different impact of PDH function on the proteins in major acetylated bands of 30 and 50 kDa proteins, coinciding with the changes in the average levels of acetylation in these bands (Figure 2). It is worth noting that the levels of phosphorylated PDHA correlate positively with succinylation of the metabolic proteins (Table 2). Because PDHA phosphorylation causes inactivation of PDH complex, this finding is in good accord with the consequences of the action of the PDH inhibitor AcPMe, that also increases succinylation of the metabolic proteins (Figure 2, the lower part). Thus, a limited decrease in PDH function, that does not change the levels of SIRT3 and SIRT5 (Figure 3), increases succinylation of the 30-50 kDa brain proteins. When the PDH inhibition increases, as occurs in the AcMeP-treated animals, the increased succinylation of 30-50 kDa proteins is overcome by increased expression of the desuccinylase SIRT5.

**Table 2.** Spearman correlations of the expression of PDHA, its Ser293-phosphorylated form (Phos-PDHA) and their ratio, with the protein acetylation and succinylation levels. The levels of the protein expression and acylation are determined by Western blots as described in “Materials and methods”. Spearman correlation coefficients Rs and statistical significance of the correlations p are shown. The statistically significant (p < 0.05) positive and negative correlations are marked in red and blue, respectively.

|  | PDHA |  | Phos-PDHA |  | Phos-PDHA / PDHA |  |
| --- | --- | --- | --- | --- | --- | --- |
|  | Rs | p | Rs | p | Rs | p |
| Ac-K Total | 0.11 | 0.64 | 0.03 | 0.90 | 0.04 | 0.87 |
| Ac-K 15 kDa | 0.03 | 0.88 | 0.02 | 0.92 | 0.01 | 0.96 |
| Ac-K 29 kDa | -0.60 | 0.00 | -0.01 | 0.95 | 0.30 | 0.18 |
| Ac-K 31 kDa | -0.52 | 0.01 | -0.17 | 0.46 | 0.15 | 0.52 |
| Ac-K 50 kDa | 0.58 | 0.00 | 0.01 | 0.97 | -0.21 | 0.35 |
| Suc-K Total | -0.02 | 0.91 | 0.47 | 0.03 | 0.25 | 0.26 |
| Suc-K 50 kDa | -0.02 | 0.94 | 0.50 | 0.02 | 0.26 | 0.23 |
| Suc-K 40 kDa | 0.06 | 0.77 | 0.44 | 0.04 | 0.24 | 0.29 |
| Suc-K 37 kDa | 0.03 | 0.89 | 0.54 | 0.01 | 0.34 | 0.12 |
| Suc-K 30 kDa | -0.14 | 0.53 | 0.27 | 0.22 | 0.13 | 0.56 |
| Suc-K 21 kDa | 0.14 | 0.54 | 0.14 | 0.53 | -0.10 | 0.67 |
| Suc-K 15 kDa | 0.01 | 0.96 | 0.15 | 0.52 | -0.10 | 0.64 |

**Figure 4.**
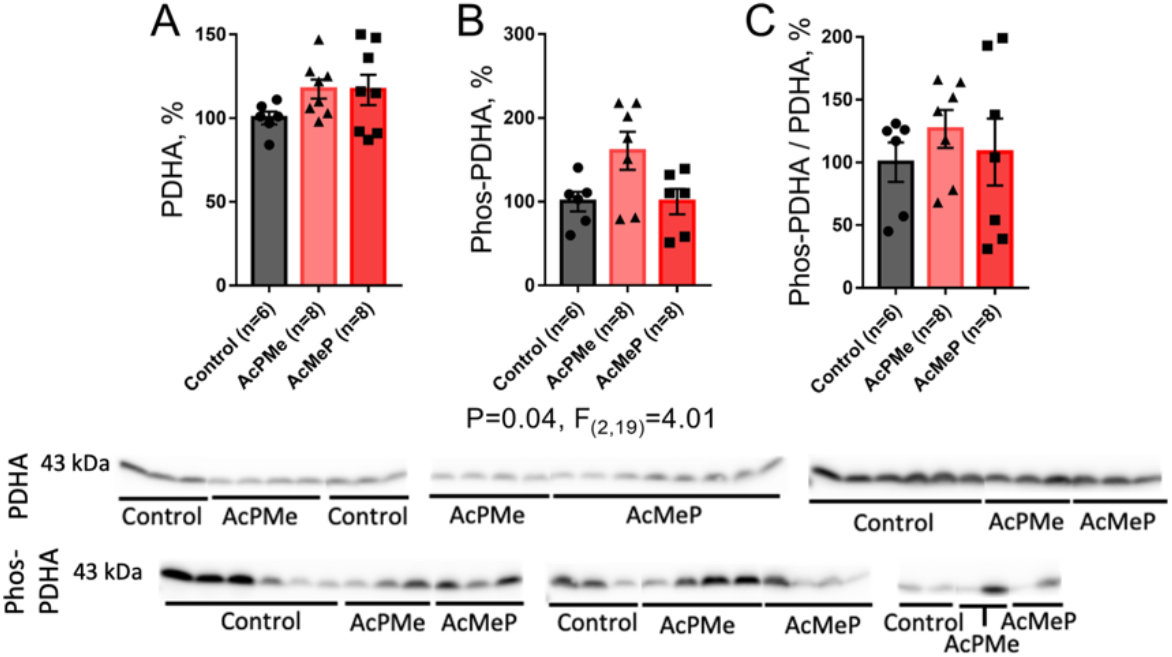
Regulation of the PDHA expression and phosphorylation in the cerebral cortex homogenates upon administration of PDH inhibitors. A – The protein levels of the phosphorylatable PDH alfa subunit (PDHA); B – the level of PDHA phosphorylated at Ser293 (Phos-PDHA); C – the ratio between the phosphorylated and total PDHA, i.e. the phosphorylation level of PDHA Ser293. PDHA and its phosphorylation are determined by Western blots shown under the graphs. Each band intensity is normalized to the total protein in the lane and related to an average value of the normalized band intensity in the control group of rats as described in “Materials and methods”. When the treatment factor is significant according to ANOVA analysis, the characteristic p and F values are given under the graphs.

### Behavioral and ECG parameters of the experimental animals do not change in response to PDH inhibitors

Physiological parameters of the AcMeP- and AcPMe-treated animals, as determined in the “Open Field” test (Figure 5, the upper part) and by non-invasive ECG (Figure 5, the lower part), do not show significant differences. This finding indicates that the biphasic biochemical changes in the brain considered above, correspond to adaptive response, i.e, provide for no deviation in the average values of physiological parameters from the control ones. It is, however, worth noting that in the absence of the significant changes in the average physiological parameters, interindividual differences in the control and treated animals may be changed by the treatments. This is exemplified by an obvious reduction in the interindividual variability of the freezing duration, accompanied by an increase in the variability of ECG parameters in the AcMeP-treated vs non-treated rats (Figure 5).

**Figure 5.**
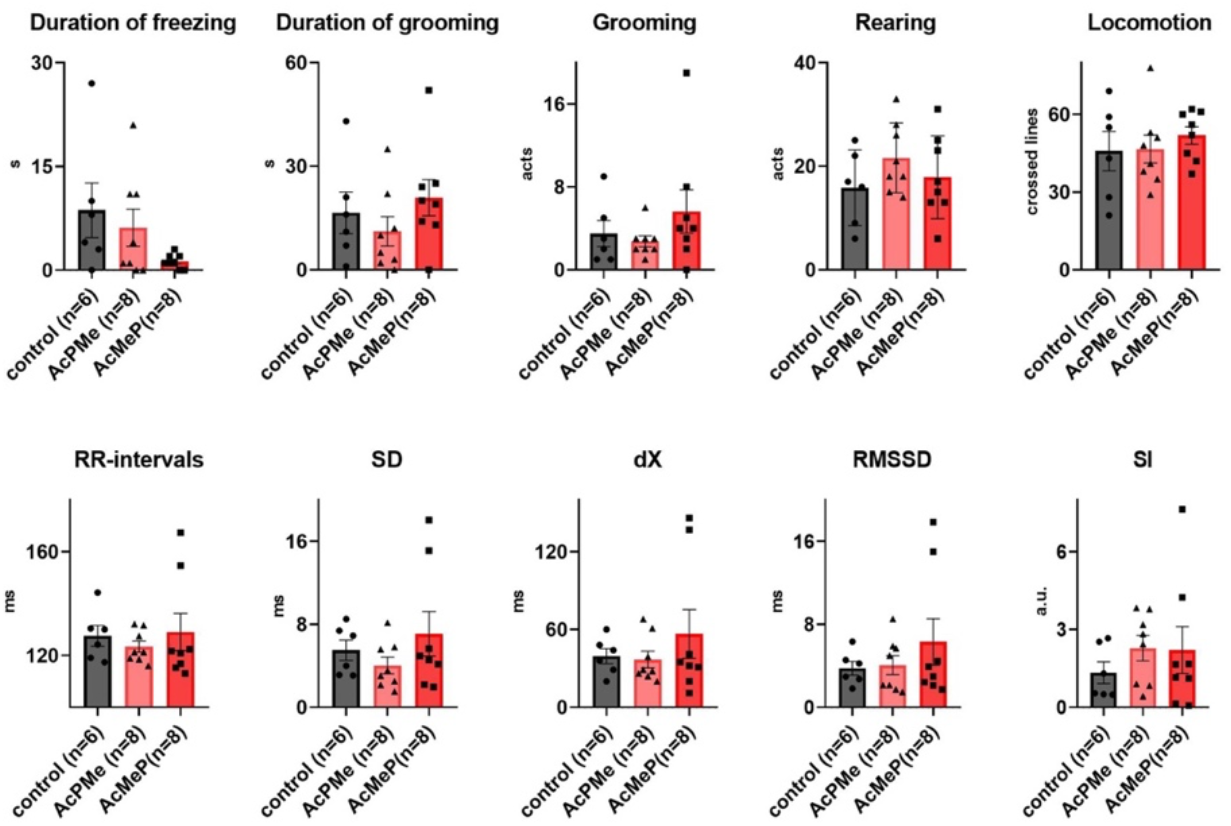
Behavioral and ECG parameters of the rats 24 hours after the administration of the PDH inhibitors. The parameters of duration of freezing and grooming, number of grooming acts, rearing acts and locomotion are obtained from the “Open Field” test as described in “Materials and methods”. The ECG parameters include the length of the RR interval, the standard deviation of the average R-R interval values (SD, ms), the heart variability rate dX, the parasympathetic (relaxation) RMSSD and sympathetic (stress) (SI) indices, estimated as described in “Materials and methods”.

Importance of the studied components of the brain acylation system for physiology is obvious from a number of significant correlations with behavioral parameters (Table 3) and with parameters of ECG (Table 4).

**Table 3.** Spearman correlations of the brain acylation system components with behavioral parameters. Spearman correlation coefficients Rs and statistical significance of the correlations p are shown. Positive or negative correlations with p < 0.1 are marked in the shades of red or blue, respectively. The statistically significant (p < 0.05) correlations are shown by the brighter colors than trends (0.05 < p < 0.1).

|  | Freezing time |  | Grooming time |  | Grooming |  | Rearing |  | Locomotion |  |
| --- | --- | --- | --- | --- | --- | --- | --- | --- | --- | --- |
|  | Rs | p | Rs | p | Rs | p | Rs | p | Rs | p |
| Sirt3 | -0.05 | 0.83 | -0.29 | 0.18 | -0.20 | 0.37 | 0.01 | 0.98 | -0.22 | 0.33 |
| Sirt5 | 0.41 | 0.06 | 0.22 | 0.33 | 0.11 | 0.61 | -0.17 | 0.44 | 0.20 | 0.37 |
| Ac-K Total | 0.12 | 0.58 | -0.06 | 0.79 | -0.49 | 0.02 | -0.03 | 0.90 | -0.26 | 0.24 |
| Ac-K 15 kDa | -0.02 | 0.94 | -0.22 | 0.33 | -0.38 | 0.08 | 0.01 | 0.97 | -0.37 | 0.09 |
| Ac-K 29 kDa | 0.07 | 0.75 | 0.14 | 0.54 | 0.05 | 0.81 | 0.19 | 0.39 | 0.24 | 0.29 |
| Ac-K 31 kDa | -0.03 | 0.88 | -0.01 | 0.97 | -0.07 | 0.76 | 0.10 | 0.67 | 0.09 | 0.69 |
| Ac-K 50 kDa | 0.12 | 0.58 | 0.29 | 0.19 | -0.26 | 0.24 | -0.16 | 0.47 | -0.16 | 0.47 |
| Suc-K Total | -0.13 | 0.56 | -0.30 | 0.18 | 0.07 | 0.76 | 0.43 | 0.05 | -0.01 | 0.97 |
| Suc-K 50 kDa | -0.17 | 0.46 | -0.17 | 0.44 | 0.15 | 0.50 | 0.31 | 0.17 | -0.04 | 0.87 |
| Suc-K 40 kDa | -0.10 | 0.67 | -0.33 | 0.13 | -0.13 | 0.55 | 0.38 | 0.08 | 0.01 | 0.96 |
| Suc-K 37 kDa | -0.25 | 0.27 | -0.35 | 0.11 | -0.03 | 0.89 | 0.30 | 0.18 | -0.18 | 0.42 |
| Suc-K 30 kDa | -0.04 | 0.85 | -0.12 | 0.59 | 0.07 | 0.77 | 0.37 | 0.09 | -0.01 | 0.96 |
| Suc-K 21 kDa | -0.27 | 0.23 | -0.29 | 0.20 | 0.11 | 0.62 | 0.41 | 0.06 | 0.06 | 0.79 |
| Suc-K 15 kDa | -0.14 | 0.52 | -0.04 | 0.87 | 0.37 | 0.09 | 0.15 | 0.50 | 0.09 | 0.68 |
| PDHA | -0.14 | 0.55 | 0.09 | 0.69 | -0.40 | 0.07 | -0.01 | 0.95 | -0.41 | 0.06 |
| Phos-PDHA | -0.12 | 0.61 | -0.14 | 0.52 | 0.07 | 0.75 | 0.00 | 0.98 | -0.16 | 0.47 |
| Phos-PDHA/PDHA | 0.05 | 0.84 | -0.21 | 0.35 | 0.11 | 0.61 | 0.04 | 0.87 | 0.11 | 0.62 |
| MDH | -0.02 | 0.94 | 0.19 | 0.40 | 0.44 | 0.04 | 0.08 | 0.74 | 0.39 | 0.07 |

**Table 4.** Spearman correlations of the brain acylation system components with ECG parameters. The parameters of ECG are deciphered in the legend to Figure 5 and “Materials and Methods”. Spearman correlation coefficients Rs and statistical significance of the correlations p are shown. Positive or negative correlations with p < 0.1 are marked in the shades of red or blue, respectively. The statistically significant (p < 0.05) correlations are shown by the brighter colors than trends (0.05 < p < 0.1).

|  | RR |  | SD |  | dX |  | RMSSD |  | SI |  |
| --- | --- | --- | --- | --- | --- | --- | --- | --- | --- | --- |
|  | Rs | p | Rs | p | Rs | p | Rs | p | Rs | p |
| Sirt3 | -0.06 | 0.79 | -0.27 | 0.23 | -0.04 | 0.87 | -0.36 | 0.10 | 0.20 | 0.38 |
| Sirt5 | -0.07 | 0.76 | -0.03 | 0.91 | -0.12 | 0.60 | -0.06 | 0.79 | 0.12 | 0.60 |
| Ac-K Total | 0.10 | 0.65 | 0.00 | 0.99 | 0.14 | 0.54 | -0.09 | 0.68 | -0.08 | 0.73 |
| Ac-K 15 kDa | 0.06 | 0.77 | -0.12 | 0.58 | 0.07 | 0.77 | -0.20 | 0.38 | 0.02 | 0.92 |
| Ac-K 29 kDa | 0.58 | 0.00 | 0.49 | 0.02 | 0.46 | 0.03 | 0.25 | 0.27 | -0.50 | 0.02 |
| Ac-K 31 kDa | 0.29 | 0.18 | 0.29 | 0.20 | 0.17 | 0.44 | 0.11 | 0.63 | -0.30 | 0.18 |
| Ac-K 50 kDa | -0.29 | 0.19 | -0.11 | 0.63 | -0.19 | 0.40 | 0.02 | 0.94 | 0.19 | 0.40 |
| Suc-K Total | 0.05 | 0.84 | -0.06 | 0.79 | 0.25 | 0.27 | 0.02 | 0.91 | -0.01 | 0.97 |
| Suc-K 50 kDa | 0.05 | 0.84 | -0.04 | 0.87 | 0.16 | 0.48 | 0.05 | 0.84 | 0.00 | 0.99 |
| Suc-K 40 kDa | 0.10 | 0.67 | -0.12 | 0.60 | 0.18 | 0.41 | -0.11 | 0.64 | 0.03 | 0.89 |
| Suc-K 37 kDa | -0.05 | 0.84 | -0.29 | 0.19 | -0.02 | 0.94 | -0.18 | 0.43 | 0.19 | 0.39 |
| Suc-K 30 kDa | 0.21 | 0.35 | -0.02 | 0.92 | 0.34 | 0.12 | 0.05 | 0.84 | -0.07 | 0.77 |
| Suc-K 21 kDa | -0.17 | 0.44 | -0.27 | 0.23 | -0.02 | 0.92 | -0.15 | 0.49 | 0.23 | 0.30 |
| Suc-K 15 kDa | -0.13 | 0.55 | -0.01 | 0.95 | 0.14 | 0.55 | 0.12 | 0.60 | 0.03 | 0.89 |
| PDHA | -0.34 | 0.12 | -0.24 | 0.27 | -0.21 | 0.34 | -0.22 | 0.33 | 0.25 | 0.26 |
| Phos-PDHA | -0.08 | 0.73 | -0.03 | 0.90 | 0.12 | 0.60 | 0.24 | 0.29 | -0.03 | 0.90 |
| Phos-PDHA/PDHA | 0.01 | 0.97 | 0.03 | 0.89 | 0.08 | 0.73 | 0.20 | 0.38 | -0.10 | 0.66 |
| MDH | -0.03 | 0.90 | 0.12 | 0.58 | 0.10 | 0.67 | -0.11 | 0.63 | -0.07 | 0.75 |

Remarkably, different components of the brain acylation system exhibit specific interdependences with physiological parameters. That is, SIRT5 levels show positive correlation with freezing time; rearing associates positively with the brain protein succinylation, while grooming acts and, to a lesser extent, locomotion are linked to the brain protein acetylation and PDHA expression (Table 3). Among the components of the brain acylation system, that correlate with the ECG parameters, acetylation of specifically 29 kDa proteins shows strong positive correlations with the parameters characterizing the heart beats frequency and variability, simultaneously negatively correlated to the sympathetic activity (stress) index (SI). The negative correlation indicates that increased acetylation of 29 kDa proteins is associated with decreased stress index. At the same time, the reversed index of parasympathetic (relaxation) activity (RMSSD) exhibits a trend to negatively correlate with the level of deacetylase SIRT3 (Table 4). This means that the higher the SIRT3 expression, the lower the relaxation. In other words, the lower level of acetylation of the brain proteins at the increased expression of SIRT3 coincides with a decreased relaxation.

## DISCUSSION

Using pharmacological inhibition of PDH by synthetic phosphonate and phosphinate analogs of pyruvate, this work has revealed specific significance of PDH function for the brain protein acylation. According to the characterized earlier [23–25] efficacy of the PDH inhibitors, administration to animals of the same doses of the inhibitors should result in different levels of the enzyme inhibition in vivo. Indeed, a hierarchical response of the brain acylation system to the PDH inhibitors is observed in this work, that is schematically summarized in Figure 6. At a moderate level (Figure 6A), the PDH inhibition decreases acetylation of 31 kDa proteins only, mostly increasing succinylation of the brain proteins. At a higher level (Figure 6B), the PDH inhibition significantly decreases acetylation of the major acetylated protein band, whose molecular mass of 15 kDa corresponds to that of histones, simultaneously increasing succinylation of these proteins. Acetylation of the proteins of other molecular masses demonstrates both decreases (29 and 31 kDa) and increases (50 kDa) at the strong PDH inhibition. However, succinylation of these non-histone proteins returns to the control value along with increased expression of desuccinylase SIRT5, compared to the state after the moderate PDH inhibition (Figure 3). As a result, succinylation of the brain proteins 30-50 kDa changes in accord with expression of SIRT5, but succinylation of the proteins of 15 kDa increases despite increased SIRT5 expression. The opposite responses of different proteins to the PDH inhibition agree with different intracellular localization of the considered proteins, such as nuclear localization of histone proteins (15 kDa) and mitochondrial/cytosolic localization of the other proteins (30-50 kDa) (Figure 6). Remarkably, in an independent study of another PDH impairment, i.e. occurring upon perturbed lipoylation of the multienzyme complex, similar opposite changes in the acetylation of different proteins are shown by using specific antibodies for histones, whose acetylation decreases, and tubulin, whose acetylation increases [37].

**Figure 6.**
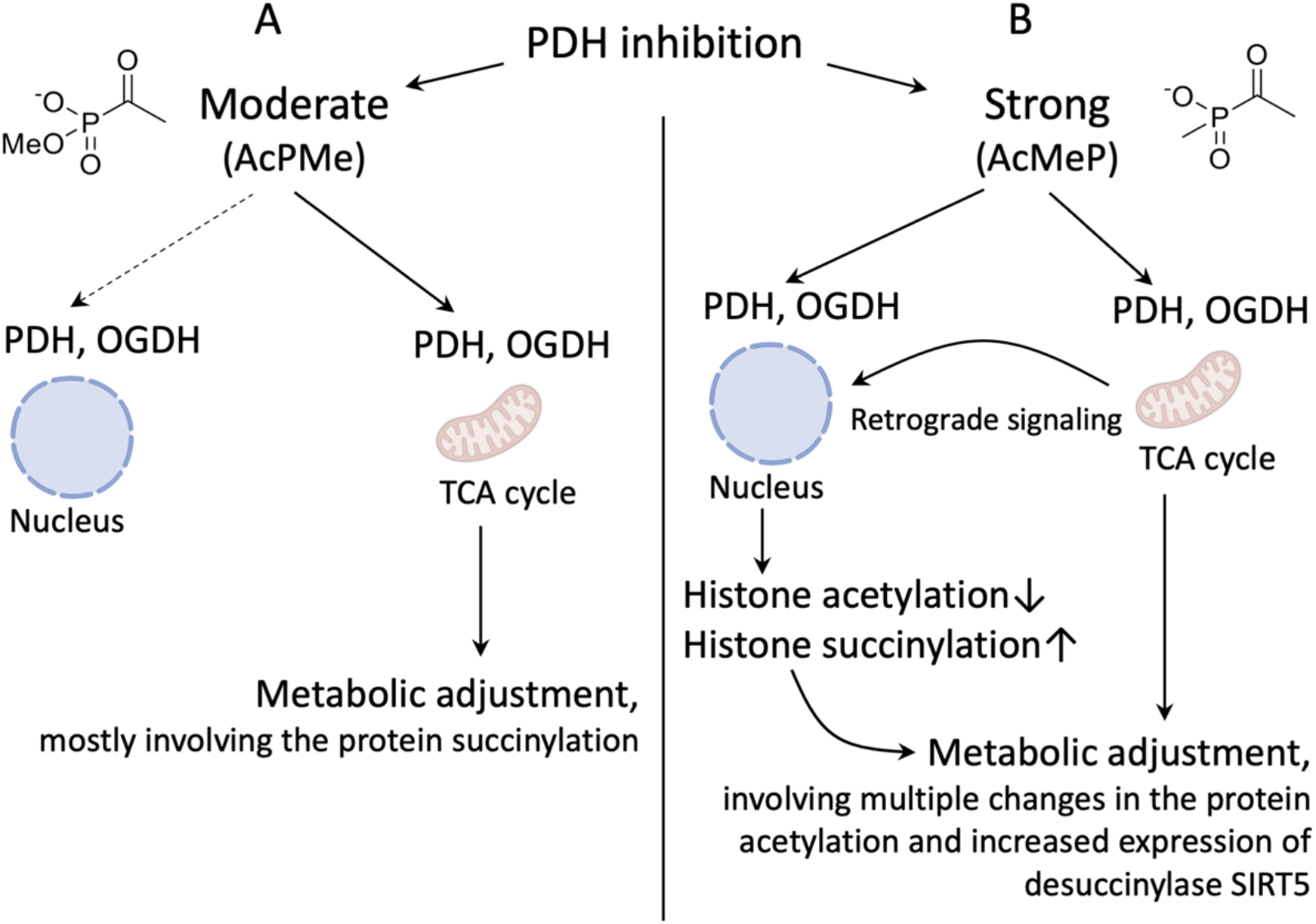
A scheme of the proposed action of the PDH inhibitors AcPMe (A) and AcMeP (B) on the metabolic and nuclear PDH and their effect for the cerebral metabolism.

It is worth noting that increased succinylation of the brain proteins of apparent molecular mass corresponding to histones (15-20 kDa) (Figure 2B) coincides with increased expression of the desuccinylase SIRT5 (Figure 3B), in good accord with activated transcription upon histones succinylation. Furthermore, the increased expression of the desuccinylase of metabolic proteins (SIRT5) decreases succinylation of the non-histone proteins (30-50 kDa, Figure 2), but not of the proteins with apparent molecular mass of histones. Remarkably, the levels of acetylation of different protein bands show strong associations with the expression of either the acetyl-CoA producer PDHA (acetylation of metabolic proteins of 30-50 kDa, Table 2) or deacetylase SIRT3 (acetylation of 15 kDa proteins, providing a major contribution into the total acetylation, Table 1). The finding indicates that the regulatory impact of the acetyl residues producer (PDH) and deacetylase (SIRT3) on the protein acetylation is not uniform, but depends on specific proteins. In particular, increased acetylation of the brain proteins with apparent molecular mass corresponding to that of histones (15 kDa), is positively correlated with the levels of mitochondrial SIRT3, suggesting a contribution of the mitochondrial protein acetylation to the retrograde signaling from mitochondria to nucleus (Figure 6). In contrast, increased expression of PDHA correlates positively with increased acetylation of proteins whose molecular mass of 50 kDa corresponds to that of tubulin. However, acetylation of the proteins of about 30 kDa correlates with the level of PDHA negatively. Thus, our data reveal a non-uniform control of the brain protein acetylation, in regards of specific components of the brain acylation system.

One feature of this complexity is a strong interdependence of different types of the protein acylations. Proteomics data stored in databases (e.g., PhosphoSite [38]) reveal that in many cases the same lysine residues may undergo different types of acylation. Due to competitive nature of such acylation reactions, it is probable that succinylation of some residues may increase when their acetylation decreases. The competition of PDH and OGDH complexes for their substrate CoA also presumes mutual effects on the corresponding acyl-CoA products from these complexes. That is, when inhibition of PDH decreases production of acetyl-CoA, increased availability of the free CoA may stimulate production of succinyl-CoA by OGDH complex. Indeed, similar to a recent study of protein acylation identified by mass spectrometry in a rat model of epilepsy [15], our current work with antibodies shows that succinylation of 30 kDa proteins positively correlates with the brain levels of SIRT3 (Table 1). The observation agrees with the competition between the acetylation and succinylation reactions due to the reasons considered above. On the other hand, both here (Table 1) and in the recent study of the rat epilepsy model [15], protein acetylation levels may correlate with the levels of the deacetylases SIRT3 and SIRT2 not negatively, but positively. This finding points to a complex interdependence of the expression of deacetylases and protein acetylation, probably due to different sources and targets of the acetylation. In contrast, in the majority of non-histone proteins the negative correlation of the protein succinylation and expression of desuccinylase of metabolic proteins (SIRT5) (Table 1) is in accordance with a straightforward view that the higher level of the desuccinylase provides for a lower level of the succinylation. Thus, competitive factors and specific regulation of the acylation sources and targets may contribute to the observed interplay of the brain protein acetylation and succinylation that we demonstrate upon the specific inhibition of PDH.

Worth noting, PDHA phosphorylation does not exhibit significant changes in response to the short-term action of PDH inhibitors. Indeed, PDHA phosphorylation is known to respond to hormonal action [39–42], that does not seem to be out of balance under our experimental conditions. As a result, adaptive response to the short-term PDH inhibition is supported by the interconnected acylation reactions rather than PDHA phosphorylation. Yet the levels of phosphorylated (inactive) PDHA positively correlate with succinylation of non-histone proteins (Table 2). Reproducing the increased succinylation upon pharmacological inhibition of PDH (Figure 2), these interdependencies of the phosphorylated (inactive) PDHA may result from the known involvement of PDHA phosphorylation into metabolic switches of the substrate fluxes between the succinyl-CoA-generating tricarboxylic acid cycle and glycolysis [43–45]. A mechanism supporting such switches may involve direct action of the phosphorylated PDHA complex with pyruvate kinase 2M as transcriptional regulator [41]. Remarkably, similar action is known for OGDH in the complex with acetyl-CoA acyltransferase 2, both controlling histones acylation by the DNA binding of the complex [12]. Study of multiple components of the brain protein acylation system reveal that memory formation and its age- or disease-dependent impairment depend not only on the long-known histone acetylation [17,46–48], but also on histone succinylation [12]. Impaired succinylation of the non-histone brain proteins has also recently been shown to be a characteristic feature of Alzheimer’s disease brains [21], where reduced OGDH activity has been a long-known hallmark [20].

Our results indicate that the coupled responses of the protein acetylation and succinylation support the stability of physiological reactions under conditions of metabolic stress. In the first line, the acylations of proteins of 30-50 kDa are changed upon PDH inhibition (Figure 6A), followed by those of proteins of 15-20 kDa (Figure 6B). We suggest that this sequence manifests the two levels of the system response to the PDH inhibition, involving first the acylation-dependent regulation of metabolic proteins and secondly the transcriptional regulation [49][50]. The correlations between the distinct components of the brain protein acylation system and physiological parameters (Table 3, 4) reveal physiological implications of the protein acylations, which are currently only emerging in the literature. In particular, both negative correlations in Table 4, i.e. that of SIRT3 with RMSSD and between the acetylation level of 29 kDa proteins and SI point to increased parasympathetic (relaxation) activity with increased acetylation of the brain proteins. This correlation may be due to the fact that the higher “acetylation potential” may manifest the acetyl-CoA-promoted biosynthesis of the neurotransmitter of parasympathetic nervous system, acetylcholine. In view of the role of the balance of the sympathetic and parasympathetic systems in controlling stress is known to be in a complex relationship with the memory formation [49], also enhancing the memory under specific conditions [50], the found link between the parasympathetic system and brain protein acetylation may contribute to the dependence of memory and cognitive functions on the brain protein acylations, discussed above. Further identification of specific proteins within the protein bands whose acylation is shown to change in response to PDH inhibitors and correlate with physiological parameters, may thus provide insights on specific components of the brain acylation system, significant for the acylation-dependent physiological functions and their impairment in diseases.

## CONCLUSIONS

Increasing inhibition of brain PDH complex decreases acetylation and increases succinylation of 15 kDa proteins. Acylations of the brain proteins of 30-50 kDa show varied responses to PDH inhibition, while PDHA phosphorylation does not change significantly. Changed acylation of the brain proteins upon PDH inhibition may not only support adaptive responses, but also be involved in pathological mechanisms.

## Author Contributions

Conceptualization, V.I.B.; validation, formal analysis, visualization, V.A.A., D.A.S., A.V.G.; writing—original draft preparation, V.I.B.; writing—review and editing, V.I.B.; all the coauthors contributed to methodology, investigation, resources and data curation; supervision, V.I.B.; project administration, V.I.B.; funding acquisition, V.I.B. All authors have read and agreed to the published version of the manuscript.

## Ethics declarations

All animal experiments were performed according to the guidelines of the Declaration of Helsinki and were approved by Bioethics Committee of Lomonosov Moscow State University (protocol number 139-a from 11 November 2021).

## Abbreviations

PDH: pyruvate dehydrogenase
PDHA: alpha subunit of PDH
OGDH: 2-oxoglutarate dehydrogenase (alpha-ketoglutarate dehydrogenase)
MDH: malate dehydrogenase
R-R interval: the average interval between the heartbeats
SI: stress index
SD: the standard deviation of the average R-R interval values
dX: the range of R-R interval values
RMSSD: the root mean square of successive differences in R-R intervals

## REFERENCES

(1) Joseph, A. D. A.; Robert, A. T.; Lawrence, R. G. Concentration-Dependent Effects of Sodium Butyrate in Chinese Hamster Cells: Cell-Cycle Progression, Inner-Histone Acetylation, Histone H1 Dephosphorylation, and Induction of an H1-Like Protein. Biochemistry 1980, 19 (12), 2656–2671. https://doi.org/10.1021/BI00553A019.

(2) Fischle, W.; Wang, Y.; Allis, C. D. Histone and Chromatin Cross-Talk. Curr Opin Cell Biol 2003, 15 (2), 172–183. https://doi.org/10.1016/S0955-0674(03)00013-9.

(3) Jeans, C.; Thelen, M. P.; Noy, A. Single Molecule Studies of Chromatin. 2006. https://doi.org/10.2172/877892.

(4) Akiyama, S. K.; Hammes, G. G. Elementary Steps in the Reaction Mechanism of the Pyruvate Dehydrogenase Multienzyme Complex from Escherichia Coli: Kinetics of Acetylation and Deacetylation. Biochemistry 1980, 19 (18), 4208–4213. https://doi.org/10.1021/BI00559A011.

(5) Waskiewicz, D. E.; Hammes, G. G. Elementary Steps in the Reaction Mechanism of the α-Ketoglutarate Dehydrogenase Multienzyme Complex from Escherichia Coli: Kinetics of Succinylation and Desuccinylation. Biochemistry 1984, 23 (14), 3136–3143. https://doi.org/10.1021/BI00309A005.

(6) Xu, Y.; Shi, Z.; Bao, L. An Expanding Repertoire of Protein Acylations. Molecular and Cellular Proteomics 2022, 21 (3). https://doi.org/10.1016/J.MCPRO.2022.100193.

(7) Bruzzone, S.; Parenti, M. D.; Grozio, A.; Ballestrero, A.; Bauer, I.; Rio, A. del; Nencioni, A. Rejuvenating Sirtuins: The Rise of a New Family of Cancer Drug Targets. ingentaconnect.com 2013, 19, 614–623.

(8) Grabowska, W.; Sikora, E.; Bielak-Zmijewska, A. Sirtuins, a Promising Target in Slowing down the Ageing Process. Biogerontology 2017, 18 (4), 447–476. https://doi.org/10.1007/S10522-017-9685-9.

(9) Sutendra, G.; Kinnaird, A.; Dromparis, P.; Paulin, R.; Stenson, T. H.; Haromy, A.; Hashimoto, K.; Zhang, N.; Flaim, E.; Michelakis, E. D. A Nuclear Pyruvate Dehydrogenase Complex Is Important for the Generation of Acetyl-CoA and Histone Acetylation. Cell 2014, 158 (1), 84–97. https://doi.org/10.1016/J.CELL.2014.04.046.

(10) Chueh, F. Y.; Leong, K. F.; Cronk, R. J.; Venkitachalam, S.; Pabich, S.; Yu, C. L. Nuclear Localization of Pyruvate Dehydrogenase Complex-E2 (PDC-E2), a Mitochondrial Enzyme, and Its Role in Signal Transducer and Activator of Transcription 5 (STAT5)-Dependent Gene Transcription. Cell Signal 2011, 23 (7), 1170–1178. https://doi.org/10.1016/J.CELLSIG.2011.03.004.

(11) Wang, Y.; Guo, Y. R.; Liu, K.; Yin, Z.; Liu, R.; Xia, Y.; Tan, L.; Yang, P.; Lee, J. H.; Li, X. J.; Hawke, D.; Zheng, Y.; Qian, X.; Lyu, J.; He, J.; Xing, D.; Tao, Y. J.; Lu, Z. KAT2A Coupled with the α-KGDH Complex Acts as a Histone H3 Succinyltransferase. Nature 2017, 552 (7684), 273–277. https://doi.org/10.1038/NATURE25003.

(12) Choi, S.; Pfleger, J.; Jeon, Y. H.; Yang, Z.; He, M.; Shin, H.; Sayed, D.; Astrof, S.; Abdellatif, M. Oxoglutarate Dehydrogenase and Acetyl-CoA Acyltransferase 2 Selectively Associate with H2A.Z-Occupied Promoters and Are Required for Histone Modifications. Biochim Biophys Acta Gene Regul Mech 2019, 1862 (10). https://doi.org/10.1016/J.BBAGRM.2019.194436.

(13) Boyko, A. I.; Karlina, I. S.; Zavileyskiy, L. G.; Aleshin, V. A.; Artiukhov, A. v; Kaehne, T.; Ksenofontov, A. L.; Ryabov, S. I.; Graf, A. v; Tramonti, A.; Bunik, V. I. Delayed Impact of 2-Oxoadipate Dehydrogenase Inhibition on the Rat Brain Metabolism Is Linked to Protein Glutarylation. frontiersin.org 2022, 9. https://doi.org/10.3389/fmed.2022.896263.

(14) Gibson, G. E.; Xu, H.; Chen, H.-L.; Chen, W.; Denton, T. T.; Zhang, S. Z. Alpha-ketoglutarate Dehydrogenase Complex-dependent Succinylation of Proteins in Neurons and Neuronal Cell Lines. Wiley Online Library 2015, 134 (1), 86–96. https://doi.org/10.1111/jnc.13096.

(15) Zavileyskiy, L. G.; Aleshin, V. A.; Kaehne, T.; Karlina, I. S.; Artiukhov, A. v; Maslova, M. v; Graf, A. v; Bunik, V. I. The Brain Protein Acylation System Responds to Seizures in the Rat Model of PTZ-Induced Epilepsy. mdpi.com 2022, 23 (20), 12302. https://doi.org/10.3390/ijms232012302.

(16) Pietrocola, F.; Galluzzi, L.; Bravo-San Pedro, J. M.; Madeo, F.; Kroemer, G. Acetyl Coenzyme A: A Central Metabolite and Second Messenger. Cell Metab 2015, 21 (6), 805–821.

(17) Mews, P.; Donahue, G.; Drake, A. M.; Luczak, V.; Abel, T.; Berger, S. L. Acetyl-CoA Synthetase Regulates Histone Acetylation and Hippocampal Memory. Nature 2017, 546 (7658), 381–386. https://doi.org/10.1038/NATURE22405.

(18) Pougovkina, O.; te Brinke, H.; Ofman, R.; van Cruchten, A. G.; Kulik, W.; Wanders, R. J. A.; Houten, S. M.; de Boer, V. C. J. Mitochondrial Protein Acetylation Is Driven by Acetyl-CoA from Fatty Acid Oxidation. Hum Mol Genet 2014, 23 (13), 3513–3522. https://doi.org/10.1093/HMG/DDU059.

(19) Gräff, J.; Tsai, L.-H. Histone Acetylation: Molecular Mnemonics on the Chromatin. Nat Rev Neurosci 2013, 14 (2), 97–111. https://doi.org/10.1038/nrn3427.

(20) Gibson, G. E.; Blass, J. P.; Beal, M. F.; Bunik, V. The α-Ketoglutarate-Dehydrogenase Complex: A Mediator between Mitochondria and Oxidative Stress in Neurodegeneration. Mol Neurobiol 2005, 31 (1–3), 43–63. https://doi.org/10.1385/MN:31:1-3:043.

(21) Yang, Y.; Tapias, V.; Acosta, D.; Xu, H.; Chen, H.; Bhawal, R.; Anderson, E. T.; Ivanova, E.; Lin, H.; Sagdullaev, B. T.; Chen, J.; Klein, W. L.; Viola, K. L.; Gandy, S.; Haroutunian, V.; Beal, M. F.; Eliezer, D.; Zhang, S.; Gibson, G. E. Altered Succinylation of Mitochondrial Proteins, APP and Tau in Alzheimer’s Disease. Nat Commun 2022, 13 (1). https://doi.org/10.1038/S41467-021-27572-2.

(22) Bunik, V. I.; Tylicki, A.; Lukashev, N. v. Thiamin Diphosphate-dependent Enzymes: From Enzymology to Metabolic Regulation, Drug Design and Disease Models. FEBS J 2013, 280 (24), 6412–6442. https://doi.org/10.1111/febs.12512.

(23) Bunik, V. I.; Artiukhov, A.; Kazantsev, A.; Goncalves, R.; Daloso, D.; Oppermann, H.; Kulakovskaya, E.; Lukashev, N.; Fernie, A.; Brand, M.; Gaunitz, F. Specific Inhibition by Synthetic Analogs of Pyruvate Reveals That the Pyruvate Dehydrogenase Reaction Is Essential for Metabolism and Viability of Glioblastoma Cells. Oncotarget 2015, 6 (37), 40036–40052. https://doi.org/10.18632/ONCOTARGET.5486.

(24) Nemeria, N. S.; Korotchkina, L. G.; Chakraborty, S.; Patel, M. S.; Jordan, F. Acetylphosphinate Is the Most Potent Mechanism-Based Substrate-like Inhibitor of Both the Human and Escherichia Coli Pyruvate Dehydrogenase Components of the Pyruvate Dehydrogenase Complex. Bioorg Chem 2006, 34 (6), 362–379. https://doi.org/10.1016/J.BIOORG.2006.09.001.

(25) Artiukhov, A. v.; Graf, A. v.; Bunik, V. I. Directed Regulation of Multienzyme Complexes of 2-Oxo Acid Dehydrogenases Using Phosphonate and Phosphinate Analogs of 2-Oxo Acids. Biochemistry (Moscow) 2016, 81 (12), 1498–1521. https://doi.org/10.1134/S0006297916120129.

(26) Bunik, V. I.; Fernie, A. R. Metabolic Control Exerted by the 2-Oxoglutarate Dehydrogenase Reaction: A Cross-Kingdom Comparison of the Crossroad between Energy Production and Nitrogen Assimilation. Biochemical Journal 2009, 422 (3), 405–421. https://doi.org/10.1042/BJ20090722.

(27) Pierozan, P.; Jernerén, F.; Ransome, Y.; Karlsson, O. The Choice of Euthanasia Method Affects Metabolic Serum Biomarkers. Basic Clin Pharmacol Toxicol 2017, 121 (2), 113–118. https://doi.org/10.1111/BCPT.12774.

(28) Suckow, M.; Stevens, K.; Wilson, R. The Laboratory Rabbit, Guinea Pig, Hamster, and Other Rodents. Academic Press 2012.

(29) Underwood, W.; Anthony, R. AVMA Guidelines for the Euthanasia of Animals: 2020 Edition. 2020, 2013 (30), 2020.

(30) Aleshin, V. A.; Graf, A. v; Artiukhov, A. v; Boyko, A. I.; Ksenofontov, A. L.; Maslova, M. v; Nogués, I.; di Salvo, M. L.; Bunik, V. I.; Belozersky, A. N. Physiological and Biochemical Markers of the Sex-Specific Sensitivity to Epileptogenic Factors, Delayed Consequences of Seizures and Their Response to Vitamins B1. mdpi.com 2021, 14 (8). https://doi.org/10.3390/ph14080737.

(31) Jackson, H. F.; Broadhurst, P. L. The Effects of Parachlorophenylalanine and Stimulus Intensity on Open-Field Test Measures in Rats. Neuropharmacology 1982, 21 (12), 1279–1282. https://doi.org/10.1016/0028-3908(82)90133-2.

(32) Liebsch, G.; Montkowski, A.; Holsboer, F.; Landgraf, R. Behavioral Profiles of Two Wistar Rat Lines Selectively Bred for High or Low Anxiety-Related Behaviour. Behavioural Brain Research 1998, 94 (2), 301–310. https://doi.org/10.1016/S0166-4328(97)00198-8.

(33) Aleshin, V. A.; Mkrtchyan, G. v; Kaehne, T.; Graf, A. v; Maslova, M. v; Bunik, V. I. Diurnal Regulation of the Function of the Rat Brain Glutamate Dehydrogenase by Acetylation and Its Dependence on Thiamine Administration. Wiley Online Library 2020, 153 (1), 80–102. https://doi.org/10.1111/jnc.14951.

(34) Tsepkova, P.; Artiukhov, A.; Boyko, A.; Aleshin, V. A.; Mkrtchyan, G. v.; Zvyagintseva, M. A.; Ryabov, S. I.; Ksenofontov, A. L.; Baratova, L. A.; Graf, A. v.; Bunik, V. I. Thiamine Induces Long-Term Changes in Amino Acid Profiles and Activities of 2-Oxoglutarate and 2-Oxoadipate Dehydrogenases in Rat Brain. Biochemistry (Moscow) 2017, 82 (6), 723–736. https://doi.org/10.1134/S0006297917060098.

(35) Ladner, C. L.; Yang, J.; Turner, R. J.; Edwards, R. A. Visible Fluorescent Detection of Proteins in Polyacrylamide Gels without Staining. Anal Biochem 2004, 326 (1), 13–20. https://doi.org/10.1016/J.AB.2003.10.047.

(36) Kim, E. Y.; Kim, W. K.; Kang, H. J.; Kim, J. H.; Chung, S. J.; Seo, Y. S.; Park, S. G.; Lee, S. C.; Bae, K. H. Acetylation of Malate Dehydrogenase 1 Promotes Adipogenic Differentiation via Activating Its Enzymatic Activity. J Lipid Res 2012, 53 (9), 1864–1876. https://doi.org/10.1194/JLR.M026567.

(37) Tong, W. H.; Maio, N.; Zhang, D. L.; Palmieri, E. M.; Ollivierre, H.; Ghosh, M. C.; McVicar, D. W.; Rouault, T. A. TLR-Activated Repression of Fe-S Cluster Biogenesis Drives a Metabolic Shift and Alters Histone and Tubulin Acetylation. Blood Adv 2018, 2 (10), 1146–1156. https://doi.org/10.1182/BLOODAD-VANCES.2018015669.

(38) Hornbeck PV, Z. B., Murray B., Kornhauser J. M., Latham V., Skrzypek E. PhosphoSitePlus, 2014: Mutations, PTMs and Recalibrations. Nucleic Acids Res 2015.

(39) Arumugam, R.; Horowitz, E.; Noland, R. C.; Lu, D.; Fleenor, D.; Freemark, M. Regulation of Islet β-Cell Pyruvate Metabolism: Interactions of Prolactin, Glucose, and Dexamethasone. Endocrinology 2010, 151 (7), 3074–3083. https://doi.org/10.1210/EN.2010-0049.

(40) Consitt, L. A.; Saxena, G.; Saneda, A.; Houmard, J. A. Age-Related Impairments in Skeletal Muscle PDH Phosphorylation and Plasma Lactate Are Indicative of Metabolic Inflexibility and the Effects of Exercise Training. Journal of Physiology-Endocrinology and Metabolism 2016, 311 (1), E145–E156. https://doi.org/10.1152/ajpendo.00452.2015.

(41) Hossain, A. J.; Islam, R.; Kim, J.-G.; Dogsom, O.; Cap, K. C.; Park, J.-B. Pyruvate Dehydrogenase A1 Phosphorylated by Insulin Associates with Pyruvate Kinase M2 and Induces LINC00273 through Histone Acetylation. Biomedicines 2022, 10 (6), 1256. https://doi.org/10.3390/BIOMEDICINES10061256.

(42) Denton, R. M.; McCormack, J. G.; Rutter, G. A.; Burnett, P.; Edgell, N. J.; Moule, S. K.; Diggle, T. A. The Hormonal Regulation of Pyruvate Dehydrogenase Complex. Adv Enzyme Regul 1996, 36, 183–198. https://doi.org/10.1016/0065-2571(95)00020-8.

(43) Kim, W.; Kaelin, J. W. G. The von Hippel–Lindau Tumor Suppressor Protein: New Insights into Oxygen Sensing and Cancer. Curr Opin Genet Dev 2003, 55–60.

(44) Li, X.; Jiang, Y.; Meisenhelder, J.; Yang, W.; Hawke, D. H.; Zheng, Y.; Xia, Y.; Aldape, K.; He, J.; Hunter, T.; Wang, L.; Lu, Z. Mitochondria-Translocated PGK1 Functions as a Protein Kinase to Coordinate Glycolysis and the TCA Cycle in Tumorigenesis. Mol Cell 2016, 61 (5), 705–719. https://doi.org/10.1016/J.MOLCEL.2016.02.009.

(45) Nie, H.; Ju, H.; Fan, J.; Shi, X.; Cheng, Y.; Cang, X.; Zheng, Z.; Duan, X.; Yi, W. O-GlcNAcylation of PGK1 Coordinates Glycolysis and TCA Cycle to Promote Tumor Growth. Nat Commun 2020, 11 (1). https://doi.org/10.1038/S41467-019-13601-8.

(46) Guan, J. S.; Haggarty, S. J.; Giacometti, E.; Dannenberg, J. H.; Joseph, N.; Gao, J.; Nieland, T. J. F.; Zhou, Y.; Wang, X.; Mazitschek, R.; Bradner, J. E.; DePinho, R. A.; Jaenisch, R.; Tsai, L. H. HDAC2 Negatively Regulates Memory Formation and Synaptic Plasticity. Nature 2009, 459 (7243), 55–60. https://doi.org/10.1038/NATURE07925.

(47) Li, X.; Zhang, J.; Li, D.; He, C.; He, K.; Xue, T.; Wan, L.; Zhang, C.; Liu, Q. Astrocytic ApoE Reprograms Neuronal Cholesterol Metabolism and Histone-Acetylation-Mediated Memory. Neuron 2021, 109 (6), 957-970.e8. https://doi.org/10.1016/J.NEURON.2021.01.005.

(48) Stilling, R. M.; Fischer, A. The Role of Histone Acetylation in Age-Associated Memory Impairment and Alzheimer’s Disease. Neurobiol Learn Mem 2011, 96 (1), 19–26. https://doi.org/10.1016/J.NLM.2011.04.002.

(49) Shields, G. S.; Sazma, M. A.; McCullough, A. M.; Yonelinas, A. P. The Effects of Acute Stress on Episodic Memory: A Meta-Analysis and Integrative Review. Psychol Bull 2017, 143 (6), 636–675. https://doi.org/10.1037/BUL0000100.

(50) Goldfarb, E. v. Enhancing Memory with Stress: Progress, Challenges, and Opportunities. Brain Cogn 2019, 133, 94–105. https://doi.org/10.1016/J.BANDC.2018.11.009.

